# Defects in exosome biogenesis are associated with sensorimotor defects in zebrafish *vps4a* mutants

**DOI:** 10.1101/2024.03.13.584854

**Authors:** Anna Shipman, Yan Gao, Desheng Liu, Shan Sun, Jingjing Zang, Peng Sun, Zoha Syed, Amol Bhagavathi, Eliot Smith, Timothy Erickson, Matthew Hill, Stephan Neuhauss, Sen-Fang Sui, Teresa Nicolson

## Abstract

Mutations in human *VPS4A* are associated with neurodevelopmental defects, including motor delays and defective muscle tone. *VPS4A* encodes a AAA-ATPase required for membrane scission, but how mutations in *VPS4A* lead to impaired control of motor function is not known. Here we identified a mutation in zebrafish *vps4a*, T248I, that affects sensorimotor transformation. In biochemical experiments we show the T248I mutation reduces the ATPase activity of Vps4a and disassembly of its substrate, ESCRT filaments, which mediate membrane scission. Consistent with the established role for Vps4a in the endocytic pathway and exosome biogenesis, *vps4a^T248I^* mutants have enlarged endosomal compartments in the CNS and decreased numbers of circulating exosomes. Resembling the central form of hypotonia in human *VPS4A* patients, motor neurons and muscle cells are unaffected in mutant zebrafish as touch sensitivity is intact. Unlike somatosensory function, optomotor responses, vestibulospinal (VS), and acoustic startle reflexes are severely impaired in *vps4a^T248I^* mutants, indicating a greater sensitivity of these circuits to the T248I mutation. ERG recordings indicate that visual ability is largely reduced in the mutants, however, *in vivo* imaging of tone-evoked responses in the inner ear and ascending auditory pathway show comparable activity. Further investigation of central pathways in *vps4a^T248I^* mutants revealed that sensory cues failed to fully activate neurons in the VS and medial longitudinal fasciculus (MLF) nuclei that directly innervate motor neurons. Our results suggest that a defect in sensorimotor transformation underlies the profound yet selective effects on motor reflexes resulting from the loss of membrane scission mediated by Vps4a.

**Significance Statement:** Here we present a T248I mutation in *vps4a*, which causes sensorimotor defects in zebrafish larvae. Spanning biochemical to systems level analyses, our study indicates that a reduction in Vps4a enzymatic activity leads to endosomal defects and reduced exosome biogenesis, resulting in pronounced deficits in vision and sensorimotor transformation of auditory and vestibular cues. We suggest that the mechanisms underlying this type of dysfunction in zebrafish may also contribute to the condition seen in human patients with *de novo* mutations in the human *VPS4A* orthologue.

## Introduction

Neurodevelopmental disorders are a collection of conditions that affect central nervous system growth and development and can have a wide variety of associated clinical signs, such as impaired cognition, difficulties in communication, and motor defects. Many neurodevelopmental disorders are the result of genetic mutations and often target molecular pathways that impact protein synthesis, transcriptional or epigenetic regulation or synaptic signaling (Lewis and Kroll, 2018; Cardoso et al., 2019; Parenti et al., 2020). An early feature that is common to several neurodevelopmental defects is central hypotonia (low muscle tone), however, it remains unclear how the underlying biochemical and sensorimotor mechanisms can result in this feature (Dan, 2022).

*De novo* heterozygous missense mutations in human Vacuolar protein sorting-associated protein 4A (*VPS4A*) have been associated with neurodevelopmental disorders characterized by pronounced delays in development, motor skills and speech, and severe intellectual disabilities and visual dysfunction (Rodger et al., 2020). Symptoms arising from problems with muscle tone such as hypotonia and dystonia are also thought to be central in nature (Ganguly et al., 2021). In addition, mutations in *VPS4A* are associated with Congenital Dyserythropoietic Anemia (Seu et al., 2020; Lunati et al., 2021). *VPS4A* encodes a type I AAA-ATPase that was originally identified in yeast. In multicellular organisms, a paralogue, *VPS4B*, is also expressed and both genes are highly conserved from plants to humans (Babst et al., 1997; Scheuring et al., 2001; Strausberg et al., 2002; Beyer et al., 2003). VPS4A binds to the <u>e</u>ndosomal <u>s</u>orting <u>c</u>omplexes required for transport (ESCRT) fission machinery to promote dissociation of the subunits via ATP hydrolysis after membrane scission occurs. Membrane scission is a key step in several cellular processes including cytokinesis, neurite severing, plasma membrane repair and multivesicular body formation (Babst et al., 2002; Monroe et al., 2014; Alonso Y Adell et al., 2016; Ott et al., 2018). Multivesicular bodies form within the endocytic pathway when late endosomes bud inward and membrane scission occurs, generating small intraluminal vesicles (Piper and Katzmann, 2007; Alonso Y Adell et al., 2016; Bebelman et al., 2020; Kalluri and LeBleu, 2020). During further processing, multivesicular bodies can subsequently either fuse to the plasma membrane to release the intraluminal vesicles as exosomes into the extracellular matrix, or fuse with lysosomes to promote degradation of its contents (Dobrowolski and De Robertis, 2012; Meldolesi, 2018). Both fates of intraluminal vesicles are key for maintaining cell homeostasis and signal transduction (Cocucci et al., 2009; Palmulli and van Niel, 2018). In animal studies, it has been previously shown that RNAi knockdown of *vps4* in fruit fly larvae results in defective pruning of neurites and knock out of *Vps4a* in mice results in lethality at preweaning stages (Loncle et al., 2015; Cacheiro et al., 2020). In line with the known function of VPS4A, fibroblasts from *VPS4A* human patients display defects in endosomal morphology and cell division (Rodger et al., 2020). How these defects lead to motor system impairment in probands is not fully understood.

Here, we identified a missense mutation in zebrafish *vps4a*, T248I, which was generated in a small scale ENU mutagenesis screen for mutants that display auditory/vestibular defects (Haffter et al., 1996). We show that the zebrafish *vps4a* mutants display endosomal abnormalities and reduced numbers of exosomes in the CNS. Both defects are consistent with the induction of stress response genes in mid- and hindbrain regions of *vps4a^T248I^* mutants.

Using *in vivo* calcium imaging, we detected strongly reduced activity in the motor circuits associated with movements evoked by sensory inputs. These findings support a model where the reduced enzymatic activity of Vps4a results in a disruption in late endosome function and exosome release, consequently reducing cellular homeostasis within neural circuits mediating sensorimotor transformation.

## Materials and Methods

### Zebrafish care and use

We maintained zebrafish lines for the *vps4a^T248I^* allele in the Top Long Fin (TLF) wild-type (WT) or *mitfa^b692^ nacre* backgrounds. The *vps4a^T248I^* Tg(α-tubulin-gal4;MP-gal4;cmlc2-GFP;UAS-GCaMP7a) line was generated by crossing into a *mitfa^b692^* Tg(α-tubulin-gal4;MP-gal4;cmlc2-GFP;UAS-GCaMP7a) fish line (a gift from Philippe Mourrain). The MP-gal4 transgene was present in most fish, marking the muscle cells as well. Breeding stocks were housed at 28.5°C and animal husbandry followed standard zebrafish methods for laboratory utilization (Westerfield, 2000), as approved and overseen by the Institutional Animal Care and Use Committees at both Oregon Health and Sciences University and Stanford University. The experiments used zebrafish larvae 3-7 days post fertilization (dpf) before gender differentiation occurs. Embryos and larvae were raised in E3 medium (0.33 mM CaCl2, 0.17 mM KCl, 0.33 mM MgSO4, and 5 mM NaCl) incubated at 28.5°C. When appropriate, larvae were anesthetized in E3 with 0.03% 3-amino benzoic acid ethylester (MESAB, Western Chemical) to minimize pain and distress.

### Genotyping Methods

For the *vps4a^T248I^* allele, we genotyped fish and larvae using kompetitive allele-specific PCR (KASP) genotyping (LGC Biosearch Technologies) per the manufacturer’s instructions.

### Identification of raumschiff mutant allele

Positional cloning was performed using simple sequence length polymorphism (SSLP) markers mapped to the zebrafish genome (Knapik et al., 1998). 5 dpf zebrafish larvae were sorted by phenotype. For Whole Genome Sequencing, 5 dpf zebrafish larvae were sorted into separate pools of 20 homozygous mutants and 20 heterozygous and homozygous wild-type siblings. Genomic DNA was collected using a DNeasy Blood & Tissue Kit (Qiagen) and samples were processed by Azenta Life Sciences (formerly GENEWIZ) for library preparation and Illumina HiSeq sequencing. The resulting .gz files were uploaded to SNPTrack (http://genetics.bwh.harvard.edu/snptrack/) to identify the mutation.

### CRISPR injections

Single guide RNAs against *vps4a* and *vps4b* were designed using the CHOPCHOP website (https://chopchop.cbu.uib.no/)(Labun et al., 2019) and synthesized (Alt-R CRISPR-Cas9 System, IDT). Three guides were chosen based on if they targeted the ATPase domain or additional exons common to all isoforms, low number of off target sites and efficiency. Multi-guide CRISPR injections were performed as described previously (Kroll et al., 2021). Briefly, crRNA and tracrRNA guides were combined in equal molar amounts in Duplex Buffer (IDT), and this RNA Duplex mix was combined with 1 μg/μL Cas9-NLS protein (PNA Bio). 1 nL of RNA guide-CAS9 mixture was injected into single cell stage TLF wildtype embryos. Embryos were checked daily for viability and auditory/vestibular phenotypes scored on 5 dpf.

### Protein expression and purification

*Saccharomyces cerevisiae* Vps4 (wild-type), and mutant Vps4(T254I) were cloned into the pET28a vector with an N-terminal His6 tag. The two proteins were expressed in Escherichia coli BL21 (DE3) strains and induced with 1 mM IPTG at 20°C overnight. Sonication was used to lyse cells and centrifuged the lysates. The supernatant was loaded into Ni–NTA beads (GE Healthcare) and the eluates were incubated with 5 U mL−1 apyrase (Sigma-Aldrich) at 4°C overnight. All proteins were further purified by SEC using a Superdex 200 10/300 GL column (GE Healthcare) with 20 mM Tis-HCl at pH 8.0, 150 mM NaCl buffer. The purity and quality of all the proteins were tested by SDS–PAGE and Nano-drop (Implen).

*Saccharomyces cerevisiae* Snf7 mutant (R52E) and Vps24 were cloned into the pET28a vector with an N-terminal His6 tag. The two proteins were expressed in Escherichia coli BL21 (DE3) strains and induced with 1 mM IPTG at 20°C overnight. Sonication was used to lyse cells and centrifuged the lysates. The supernatant was loaded to a Ni–NTA beads (GE Healthcare) and the elute was further purified by SEC using a Superdex 200 10/300 GL column (GE Healthcare) with 20 mM Tis-HCl at pH 8.0, 150 mM NaCl buffer. The purity and quality of all the proteins were tested by SDS–PAGE and Nano-drop (Implen).

*Saccharomyces cerevisiae* Vps2 were cloned into the pET28a vector with an N-terminal MBP-His6 -TEV-tag. We expressed the resulting Vps2 construct in E.coli BL21 (DE3) at 20°C for 16 hours with 1 mM IPTG. We purified the protein using Ni–NTA resin (GE Healthcare). We mixed the eluted fractions from the His column with TEV (tobacco etch virus) protease for 3 hours at 4°C. We then loaded the cleaved samples onto a 5 mL HiTrap Q-sepharose FF column (GE Healthcare) and eluted the material with a gradient of 0%-100% buffer (20 mM Tris-HCl, pH 8.0, 500 mM NaCl) in 50 column volumes.

### Sedimentation Analysis of Filament Disassembly

Yeast ESCRT III filaments were assembled by mixing Snd7(R52E), Vps24 and Vps2 in a molar ratio of 2:1:1 at room temperature for 1 hour. Then, the formed filaments were centrifuged at 50,000 r.p.m. for 30 minutes in a TLA-100 ultracentrifuge rotor (Beckman Coulter). The pellet was resuspended in the buffer containing 20 mM Tris-HCl at pH 8.0, 150 mM NaCl. The pellet (P) and supernatant (S) were analyzed by SDS–PAGE.

For sedimentation analysis of ESCRT III filament disassembly, filaments were mixed with 5 μM Vps4 or Vps4(T254I) in 20 mM Tris-HCl at pH 8.0, 150 mM NaCl, 2 mM MgCl2 buffer, separately. Reactions were started by adding ATP to a final concentration of 5 mM. After incubation at room temperature for 10 minutes, reactions were stopped by adding EDTA to a final concentration of 50 mM. Samples were then subjected to ultracentrifugation and analyzed via SDS-PAGE and negative staining.

### ATPase activity assay

ATPase activities were determined for wild-type and mutant Vps4 proteins using the Quantichrome ATPase/GTPase Assay Kit (BioAssay Systems) at room temperature. Briefly, 5 μM Vps4 or mutant Vps4(T254I) protein was added to ATPase assay buffer and 4 mM ATP for 30 minutes, and then stopped using reagent from the kit. The solution was incubated at room temperature for 30 minutes, and immediately the absorbance at 620 nm was detected using a plate reader (Enspire, Perkin Elmer). The released phosphate was calculated based on the absorbance standard curve established using KH2PO4 standards.

### Auditory-evoked behavioral response (AEBR)

AEBR was tested as described previously (Erickson et al., 2017; Smith et al., 2020) using the Zebrabox system (ViewPoint Life Sciences). 5 dpf wild-type siblings or *vps4a^T248I^* mutant larvae were confined to wells of a 96-well plate in 200 μL E3 per well, in the dark. Larvae were subjected to 100 millisecond, 1kHz pure tones at 157 decibel sound pressure (dB SPL, relative to 1mPA) every 2 seconds for 1 minute. An infrared camera recorded larval movements and ZebraLab tracking software documented pixel changes, which represent movement over time for each larva. To distinguish between evoked responses and spontaneous background movement, movements within 1 second before a stimulus were excluded from a given trial. For each larva, the positive responses were calculated as a percentage out of total possible response within a minute, and the best of three trials was used for analysis.

### Vestibulospinal Reflex (VSR)

The device and method used to test the VSR was done as previously described (Sun et al., 2018; Gao and Nicolson, 2021). Briefly, 5 dpf larvae were embedded dorsal-up in 2% prewarmed low melt agarose in E3 media without anesthetic on a mounting chamber. The agarose around the trunk and pectoral fins was carefully removed and replaced with 20 μL E3 media, and the mounting chamber was placed in the device. A motor rocks the platform from −75° to + 75° at 0.53 Hz, and an infrared camera recorded video of the reflex at 30 fps (1280 frames).

Modified ZebraZoom software (Mirat et al., 2013) tracks the movements of the trunk at regular intervals and calculates the real tail angle over time between the X axis and the axis formed by the tip of the tail and the center of the swimming bladder.

### Vestibular-induced eye movement (VIEM)

The VIEM experiment was done similarly as described previously using a modified device (Mo et al., 2010; Sun et al., 2018). 5 dpf larvae were embedded on a small cover slip in prewarmed 2% low melt agarose in E3 buffer, without anesthetic. Agarose surrounding the eyes was gently removed and replaced with approximately 10 μL of E3 media to permit free eye moment. The mounted larvae were positioned vertically, head down, between an infrared camera and an infrared LED array on a programmable rocking platform. Larvae kept in darkness during the experiment. The platform rocked (-45° to 45°) at 0.25 Hz frequency for 60 seconds while larval eye movements were video recorded in infrared.

Software-based video analyses quantified the amplitudes of angular eye movements as a function of their frequency of repetition. Larvae with normal VIEM show peak movement amplitude at 0.25 Hz, in synchrony with platform rotation. Baseline spontaneous eye movement, taken as the average amplitude across all quantified frequencies, was subtracted to determine the VIEM amplitude at 0.25 Hz. We calculated the normalized amplitude of the eye ratio for each larva and used the two-tailed unpaired t test with Welch’s correction to compare mutants to their corresponding siblings.

### Touch Startle Reflex

Touch Startle was performed by mounting 7 dpf zebrafish larvae similarly to the VSR mounting method. A micromanipulator holding a borosilicate glass capillary tube (Sutter, #BF150-86-10) pulled into a needle was used to touch the head of the larvae. Startle movements were recorded by a Hamamatsu C11440 ORCA-Flash 2.8 camera at 100 fps. Quantification of C-bends was performed as described in (Budick and O’Malley, 2000).

### Optomotor Reflex (OMR)

To test for the optomotor reflex, we adapted the assay from Neuhauss et al., 1999. A set of 4 tracks (each lane measuring at 12 cm x 0.6 cm) and comb that fit between the lanes, was constructed using 3D printing. The lanes were placed in a 15 cm petri dish (VWR) and filled partially with E3 buffer. The comb was inserted to block the starting end of the lanes, and 5-6 dpf larvae were placed in the starting area of the lanes. The petri dish was placed over a screen that plays a video of 0.5 cm stripes. To record the movement of the larva along the lanes, a camera was mounted above the petri dish. To perform the experiment, the video of stripes was played, and the comb was removed, allowing larva to swim against the movement of stripes to the end of the lane. To quantify this reflex, screenshots of the video were taken of the larva in the starting region of the lane prior to comb removal and then 30 seconds after the comb was removed. The distance between the location of each larva prior to comb removal and 30 seconds after was measured in ImageJ in millimeters.

### Electroretinogram (ERG)

ERG recordings were performed on 5 - 6 dpf mutant and wild-type sibling larvae, as described in (Makhankov et al., 2004; Niklaus et al., 2017). b-wave amplitudes were plotted and analyzed using Dunn’s multiple comparisons.

### Immunohistochemistry

Larvae were fixed with 4% PFA, dehydrated with Methanol, rehydrated using a graded series of PBSDT (1x PBS with 0.25% Tween-20 and 1% DMSO): Methanol and then treated with 150 mM Tris-HCl for antigen retrieval. Samples were then washed and permeabilized with ice cold 100% acetone, followed by more washes. Samples were blocked for 2 hours at room temperature with 6% goat serum and 3% BSA, then incubated overnight with 1:1000 anti-acetylated tubulin rabbit primary antibodies (ThermoFisher). Samples were then incubated overnight with 1:1000 anti-rabbit Alexa 488 (Invitrogen) secondary antibodies, protected from light.

### Lysotracker

5 dpf PTU-treated larvae were sorted and placed in mutant-sibling pairs in a 2 mL Eppendorf tube. LysoTracker Red DND-99 (Invitrogen) was diluted to 1:100 to examine brains or 1:200 to examine inner ear hair cells. Larvae were incubated in Lysotracker for 40 minutes at 28.5°C with the tube left open but protected from light. The larvae were then washed with E3 buffer for 10 minutes and immediately imaged.

Hindbrain lysotracker uptake was quantified by selecting a single Z-stack slice of the hindbrain region and thresholding it to highlight particles in the size range of endosomal compartments (size = 0.02-2 μm). The integrated density and area of particles from two equally sized regions of interest from hindbrain sections and a single region of neuropil were measured from each slice using the Analyze Particles function of ImageJ. The integrated density measurements were averaged and normalized to the integrated density of the neuropil.

### CD63-phluorin Injections and Analysis

pUbi-CD63-pHluorin was a gift from Frederik Verweij and was generated as described in (Verweij et al., 2019). Nacre-background *vps4a^T248I^* mutant and sibling embryos were injected at the one cell stage with 100 ng/μL pUbi-CD63-pHluorin. Damaged embryos were sorted out, and only larvae with CD63-pHluorin expression on muscle cell membranes were selected for live imaging on 3-4 dpf. Time lapse videos were recorded at 50 fps for 2 minutes at 63x with a 2x optical zoom. For analysis, three frames were combined, and the mean pixel intensity was measured from regions of the ventricular intracranial space, surrounding tissue as background, and muscle cells. The mean pixel intensity of ventricular intracranial space was normalized to background and then divided by muscle cell pixel intensity.

### Bulk Tissue RNASeq

Total RNA was isolated from 5 dpf wild type and vps4a mutant larvae (n = 15-20 each genotype) using TRIzol reagent (Thermo Fisher, 15596026) according to the manufacturer’s protocol. RNA-seq library construction and single-end sequencing were performed by the OHSU Massively Parallel Sequencing Shared Resource (RRID SCR_009984, Oregon Health & Science University, Portland, OR, USA). Sequencing reads were mapped to the Danio rerio GRCz10 genome using Tophat v2.1.1 (Kim et al., 2013) from within an OSHU instance of the Galaxy sequence analysis platform (Afgan et al., 2016). The resulting BAM files were imported to SeqMonk v1.42.0 (www.bioinformatics.babraham.ac.uk/projects/seqmonk/) for data visualization and statistical analysis using the Intensity Difference filter for unreplicated data sets. Genes with an adjusted p-value < 0.05 were considered differentially expressed.

### Hybridization Chain Reaction (HCR)

For HCR, custom probes for *vps4a*, *atf3*, *jun*, *gap43, opn1sw1* and the corresponding fluorescent B-amplifier hairpins were ordered from Molecular Instruments (https://www.molecularinstruments.com/). Probes were chosen based on their log2(fold change) and number of reads from bulk RNA sequencing analysis. Probes were assigned B-amplifiers with the intention of doing multi-plex experiments. Probe sequences are considered confidential and proprietary information of Molecular Instruments based on their Terms and Conditions, so further inquiries may be directed to their technical team.

PTU-treated larvae were sorted at 5 dpf and fixed overnight in 4% paraformaldehyde (PFA) at 4°C with rocking. After fixation, larvae were washed in PBST (1X PBS with 0.1% Tween-20), dehydrated with a methanol:PBS series (25%, 50% and 75% methanol in PBS), then stored at least overnight at -20°C. Larvae were subsequently rehydrated with a reverse methanol:PBST series, washed several times with PBST, permeabilized for 15 minutes with 20 μg/mL Proteinase K, post-fixed with 4% PFA and then washed again. Probe detection and amplification with fluorescent hairpins was performed as recommended by the MI protocol for whole mount zebrafish larva with extended pre-hybridization and pre-amplification times (4 hours each) and probe concentrations adjusted based on the reads of each probe (2 nM for *atf3* and *jun*, 5 nM for *gap43*, 10 nM for *vps4a*).

### Calcium Imaging

Larvae at 3 dpf *vps4a^T248I^ Tg(α-tubulin-gal4:UAS-GCaMP7a)* were sorted for strong GCaMP expression in the brain. 5 dpf GCaMP positive larvae were mounted in 2% low melt agarose on a custom depression slide fitted with a mini-speaker (DAEX-9-45M Skinny Mini Exciter 9 mm 1 W 4 Ohm). Using an AIYIMA A07 amplifier, tone stimuli were played through the mini speaker using a custom script in MATLAB (R2020b, MathWorks). 600 Hz (0.35 V) or 450 Hz (0.2 V) tones were played for 0.6 seconds or 0.1 seconds, respectively, five times with 10 second intervals. Time lapse images and fluorescence values were recorded using LAS-X software (Ver 3.7.2.22383). To account for variability in GCaMP7a expression between larvae, all fluorescence values were normalized by subtracting background measurements. Traces were generated by averaging the normalized fluorescence value (F) for each time point across all larvae and plotting this value over time. To calculate percentage change in fluorescence (ΔFt/F0%), 2-3 frames, containing image artifacts (due to the tone stimulation shaking the slide) were identified for each stimulus, and fluorescence values corresponding to these frames were discarded. The F0 measurement was selected from the frame prior to tone stimulation, based on the time lapse video and presence of artifact frames, and Ft was selected as the highest frame immediately after tone stimulation. (ΔFt/F0)/F0 was used to calculate the change in GCaMP7a fluorescence and was multiplied by 100 to convert into a percentage to generate “(ΔFt/F0%)”. The average percentage change in fluorescence was then calculated for each larva and differences between wild-type siblings and *vps4a* mutants was analyzed using unpaired t tests.

### Retrograde Labeling of Mauthner Cells

Retrograde labeling was adapted from previously published methods (Gahtan et al., 2002; Wehner et al., 2017). Larvae were anesthetized with 0.3% MESAB prior to spinal injection of 10 mg/mL 10,000 MW Tetramethylrhodamine dextran (10,000 MW, D1866, Thermo Fisher) in PBS. Custom glass injection needles were prepared from borosilicate glass capillaries (TW100F-4, World Precision Instruments) using a micro-needle puller (Suttner Instruments Co, Model P-87) using heat cycle 517, pulling cycle 75, velocity 75 and time 90, and then the tip was broken using forceps. Larvae were mounted laterally in 2% low melt agarose for injection, and then incubated in E3 media overnight to recover before imaging.

### Microscopy and Image Processing

Images for LysoTracker, CD63pHluorin, and HCR were take using an upright LSM700 laser-scanning confocal microscope with Zeiss Zen software (Carl Zeiss). For live imaging of larva, they were anesthetized in 0.03% MESAB in E3 media and mounted in 2% prewarmed low melt agarose. Images were collected using 488 nm, 555 nm and 647 nm lasers, depending on the requirements of the individual experiment (Acetylated Tubulin and CD63pHluorin used 488 nm, LysoTracker used 555 nm, HCR used up to all three depending on the fluorescent hairpins used). We used a Leica HC PL Fluotar 10x/0.3 lens for Acetylated Tubulin, HCR and LysoTracker images, a Plan Apochromat 40x/1.0DIC lens for LysoTracker quantification images and an Acroplan 63x/0.95W water immersion lens for higher magnification HCR images. Laser power and gain were determined independently for each fluorophore and kept consistent within each experiment. The FIJI build of ImageJ Version 2.1.0/1.53t was used to process all raw image files. For Calcium Signaling, scans were taken using a Leica SP-8 Confocal Microscope equipped with a 20x (NA 1.0) objective, using LAS-X software. Images were collected using 488 nm white laser light.

### Experimental Design and Statistical Analysis

The number of zebrafish larvae used is denoted in the figure legends. G*Power software was used to ensure there were sufficient samples tested. All statistical analyses were performed using GraphPad Prism software (version 9). The data was tested for normal distribution using Shapiro-Wilk test. Data is presented as mean ± SD, and significance between mutant and wild-type sibling groups was determined using unpaired t test unless otherwise noted. The image in figure 1C was adapted from Sun et al., 2017 using Mol* (Sehnal et al., 2021) and RCSB PDB (PDB ID:5XMI) software.

**Figure 1.**
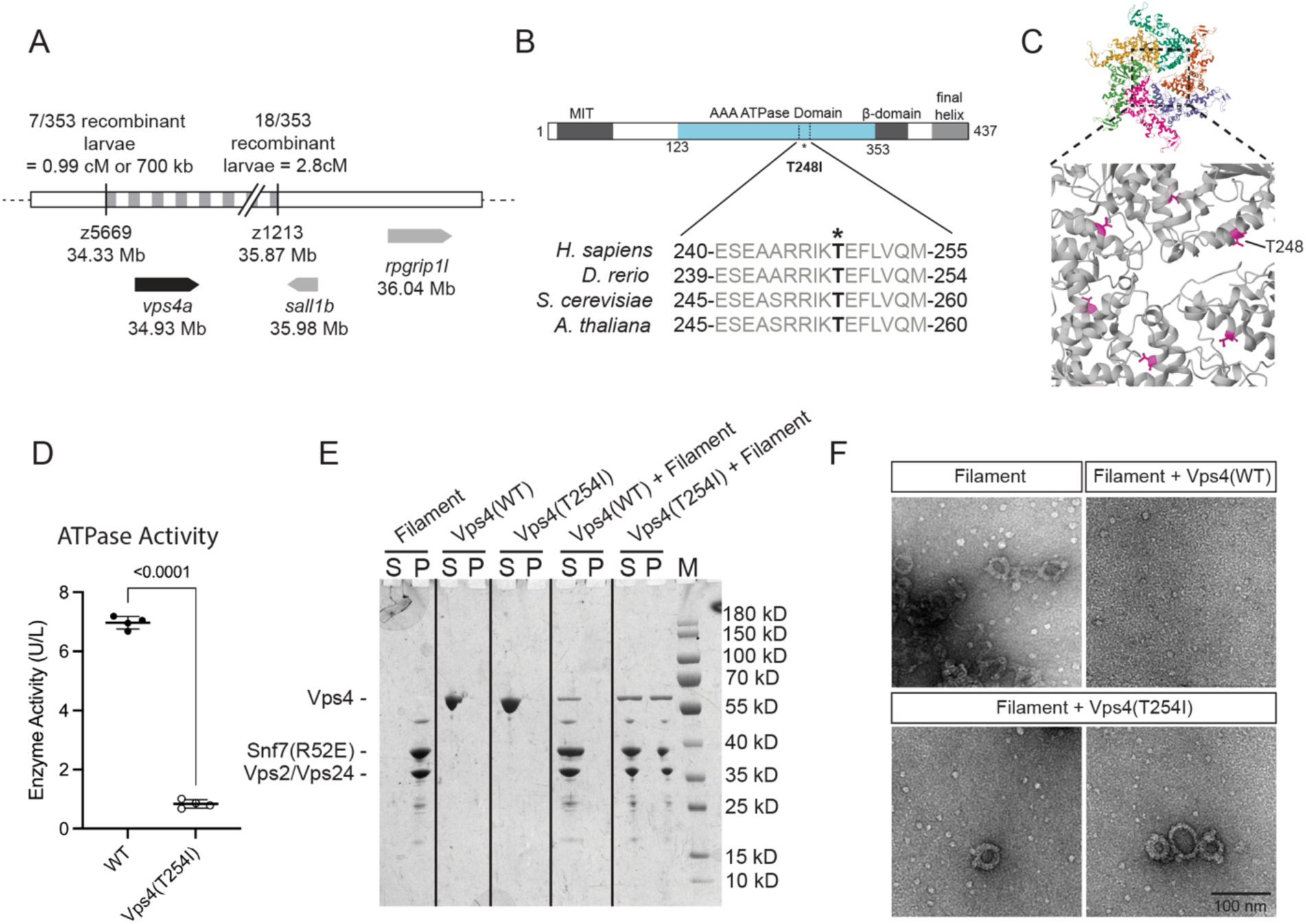
The T248I mutation in *vps4a* reduces ATPase activity and ESCRT-III filament disassembly. **A,** *vps4a* is located on Chromosome 25 in a location corresponding to where classical mapping methods have identified the location of the gene responsible for the *raumschiff* phenotypes. **B,** Bar diagram of Vps4a protein showing the location of the T248I mutation. **C,** Atomic model structure of the Vps4^E233Q^ hexamer, with T248 marked in magenta (Image adapted from the RCSB PDB ID:5XMI, using Mol*). **D,** Mutant Vps4 (T254I) yeast protein has decreased ATP activity compared to wild-type Vps4 protein. Unpaired t test, *p* < 0.0001, nWT = 4, n*vps4(T254I)* = 4, *t* = 47.92, df = 6. **E,** Sedimentation analysis of Vps4 and Vps4(T254I)-mediated disassembly of ESCRT-III filaments. Pellet (P) and supernatant (S) fractions were analyzed by SDS-PAGE. Marker (M). **F,** Negative staining EM micrograph of sedimentation analysis of ESCRT-III filament disassembly.

## Results

### A missense mutation in *vps4a* reduces ATPase activity and ESCRT-III filament disassembly

*raumschiff* mutants were generated from an ENU mutagenesis screen for genes that affect larval auditory and vestibular function (Haffter et al., 1996). Mutant larvae fail to respond to auditory cues and maintain an upright posture. The mutation is recessive and fully penetrant at 5 days postfertilization (dpf) with lethality occurring around 9-10 dpf. To identify the causative gene of the *raumschiff* phenotype, we used whole-genome sequencing and identified a threonine to isoleucine missense mutation in *vps4a* using SNPTrack (Leshchiner et al., 2012). The critical interval for *raumschiff* was also mapped to a region approximately 700 kb (or 1 cM) south of the polymorphic marker z5669, placing the locus at 34.33 Mb on Chromosome 25. *vps4a* is located at 34.93Mb on Chromosome 25, which is within the predicted region for the causative gene. No other polymorphisms were identified by SNPTrack in this region (Fig. 1A).

Vps4a is a hexameric type I AAA-ATPase member of the ESCRT-III complex, whose role is to promote the dissociation of the ESCRT-III subunits via ATP hydrolysis (Babst et al., 1997; Davies et al., 2010; Monroe et al., 2014; McCullough et al., 2018). The threonine to isoleucine mutation (T248I) we identified is present in the AAA ATPase domain in a region that is highly conserved between species (Fig. 1B). To determine whether the T248I mutation was comparable to a nonsense allele of *vps4a*, we edited the *vps4a* locus using multiple CRISPR/Cas9 guides (Wu et al., 2018; Kroll et al., 2021). In our experiments, multi-guide injections resulted in a G0 phenotype in 16.9% of injected larvae that was indistinguishable from the auditory/vestibular phenotype observed in *vps4a^T248I^* mutants (400 embryos were injected at < 1 hour past fertilization; of the larvae that survived to 5 dpf, 56/331 larvae showed absence of acoustic startle reflexes and abnormal posture). These results suggest that the T248I mutation is comparable to a null allele of Vps4a. In addition, the *vps4b* paralogue is also ubiquitously expressed (Yang et al., 2016), therefore we generated *vps4a/vps4b* G0 double mutants using CRISPR-based editing to test whether *vps4b* could compensate for the loss of *vps4a* during development. A previous study using morpholinos against *vps4b* reported a tooth phenotype analogous to human dentin dysplasia I (Yang et al., 2016). We observed that embryos injected with multiple CRISPR guides against *vps4b* alone, or *vps4a* and *vps4b* together, developed normally until 3 dpf. The lack of embryonic lethality is presumably due to the presence of maternal RNA during early developmental stages. However, in contrast to the morpholino study, *vps4b* and *vps4a/4b* multi-guide injected larvae exhibited progressive necrosis and by 5 dpf, there was complete lethality (558 embryos were injected with *vps4b* guides; 468 larvae survived to 1 dpf, and 2 survived to 5 dpf. 298 embryos were injected with *vps4a/4b* guides; 236 larvae survived to 1 dpf, and 1 survived to 5 dpf). Our results suggest that *vps4b* is required for global cellular homeostasis whereas *vps4a* selectively affects the nervous system. A similar effect was reported for the Nsfa and Nsfb AAA-ATPases that mediate vesicle fusion; mutations in zebrafish *nsfa* largely affect the nervous system, while the loss of *nsfb* functions leads to general necrosis at 4 dpf (Woods et al., 2006; Hanovice et al., 2015; Gao et al., 2023).

Based on the Cryo-EM structure of the yeast Vps4 AAA-ATPase (Sun et al., 2017), threonine 248 is located at the interface between Vps4a hexamer subunits (Fig. 1C). To assess the effect of the isoleucine substitution on ATPase activity, we generated an equivalent mutation, T254I, in the AAA ATPase domain of yeast Vps4. When tested for ATPase activity *in vitro*, Vps4^T254I^ had severely reduced enzymatic activity compared to wild-type Vps4a, although some activity remained (*p* < 0.0001, nWT = 4, n*vps4(T254I)* = 4, *t* = 47.92, df = 6) (Fig. 1D).

To determine whether the residual function of the T254I protein could mediate filament disassembly, we incubated Vps4^T254I^ with the ESCRT-III filaments formed by Vps2, Vps24 and Snf7 *in vitro* and used sedimentation to assess disassembly. In our assay, disassembled ESCRT-III filament proteins are present in the supernatant whereas assembled filaments sediment to the pellet fraction (Sun et al., 2017). We observed that wild-type Vps4a completely disassembled the ESCRT-III filaments, with all proteins found in the supernatant (Fig. 1E). In contrast, mutant Vps4^T254I^ only partially disassembled the ESCRT filaments (Fig. 1E). Using negative stain electron microscopy (EM), assembled ESCRT-III filaments appear as rings or have corkscrew structures (Fig. 1F, upper left panel). When incubated with wild-type Vps4, filament structures were absent, indicating full disassembly (Fig. 1F, upper right panel). In the presence of Vps4^T254I^, assembled filaments were still detectable albeit there were fewer filaments compared to samples where no Vps4 protein was added (Fig. 1F, lower panels).

These results demonstrate that the T245I mutation at the interface between Vps4a subunits reduces but does not fully abolish the ATPase activity of the protein, and this reduction in activity is reflected in the incomplete disassociation of ESCRT-III filaments.

### The *vps4a^T248I^* mutation results in enlarged endosomal membrane compartments and a reduction of circulating exosomes in the CNS

Using RNA-FISH, we found that the expression of *vps4a* transcripts is ubiquitous at 5 dpf, with higher levels expressed in the CNS and retina (Fig. 2A). In light of the higher expression of *vps4a* in the CNS and the neurological disorder in human patients, we focused our attention on the larval brain. As stated previously, Vps4a mediates the formation of multivesicular bodies from late endosomes. To determine if the mutation affects this function in the nervous system, we performed *in vivo* imaging of LysoTracker, a vital dye that accumulates in acidic membrane compartments such as low pH endosomal compartments. As shown in Fig. 2B, endocytic compartments in *vps4a^T248I^* mutants labeled more brightly with LysoTracker compared to wild-type siblings. The average size and pixel density of acidic membrane compartments in the hindbrain were quantified and we found that *vps4a^T248I^* mutants have significantly larger acidic membrane compartments (*p* = 0.0015, nWT = 7, n*vps4a(T248I)* = 10, *t* = 3.873, df = 15) (Fig. 2C, left panel) that take up more dye (*p* = 0.0020, nWT = 7, n*vps4a(T248I)* = 10, *t* = 3.744, df = 15) (Fig. 2C, right panel). Together, these results imply that the *vps4a^T248I^*mutation has a detrimental effect on endocytic compartment processing in the CNS.

**Figure 2.**
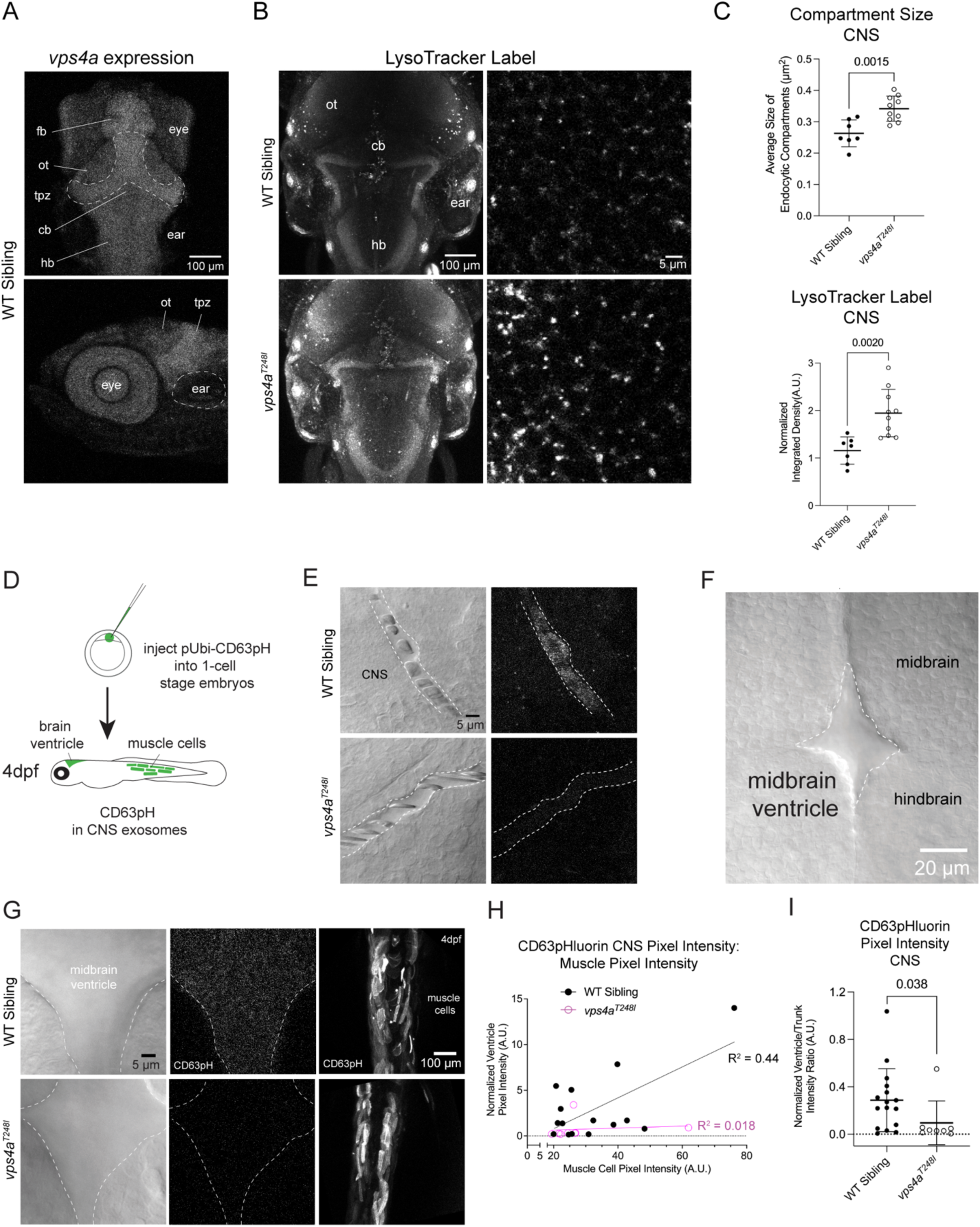
*vps4a^T248I^*mutants have enlarged endosomal compartments and release fewer exosomes in the CNS. **A,** Confocal Z-projections of a 5 dpf wild-type zebrafish larva labeled with HCR RNA-FISH probes for *vps4a.* fb, forebrain; ot, optic tectum; tpz, tectal proliferation zone, cb, cerebellum; hb, hindbrain. **B,** LysoTracker staining of 5 dpf wild-type siblings and *vps4a^T248I^* mutants show an increased staining in the mutant. **C,** Acidic membrane compartment size in the hindbrain of *vps4a^T248I^* mutants (N = 10) is significantly larger than those in wild-type siblings (N = 7); unpaired t test, *p* = 0.0015, *t* = 3.873, df = 15. LysoTracker vital dye taken up by *vps4a^T248I^*mutants (N = 10), measured as normalized pixel integrated density, is significantly higher compared to wild-type siblings (N =7); unpaired t test, *p* = 0.0020, *t* = 3.744, df = 15. **D,** Schematic of injecting 1-cell stage nacre-background embryos with pUbi:CD63-pHluorin. Healthy larvae were sorted on 1 dpf, and larvae expressing CD63-pHluorin in muscle cells was sorted on 3 dpf, for imaging on 4 dpf. **E,** Decreased CD63-pHluorin expression in 3 dpf *vps4a^T248I^* mutant CNS blood vessels. **F,** Dorsal view of 4 dpf zebrafish larva showing the midbrain ventricle. **G,** Decreased CD63-pHluorin expression in *vps4a^T248I^* mutant brain ventricles, but trunk muscles express comparable levels to wild-type siblings. **H,** Pixel intensity of muscles cells is correlated with normalized pixel intensity of CD63-pHluorin signal in the midbrain ventricle in wild-type siblings (N = 16). **I,** Quantification of CD63-pHluorin fluorescence in wild-type siblings (N = 16) and *vps4a^T248I^* mutants (N = 8); Mann-Whitney test, *p* = 0.0382 (exact).

Given the known role of Vps4a in multivesicular body formation, we examined whether the mutation affects the number of exosomes that are released upon fusion of multivesicular bodies to the plasma membrane. To detect circulating exosomes in live fish, we used CD63-pHluorin, a pH-sensitive GFP reporter that is quenched in intraluminal vesicles in low pH multivesicular bodies and fluoresces when these vesicles are released into a higher pH extracellular milieu as exosomes (Verweij et al., 2018). We microinjected a plasmid for ubiquitous expression of CD63-pHluorin into single-cell stage embryos, and selected for larvae that transiently expressed the CD63-pHluorin based on signal in the muscle cells, which is readily detectable by 3 dpf (Fig. 2D). The plasma membrane of muscle cells was commonly labeled in our injected fish as was previously reported (Verweij et al., 2018) and served as a positive control for expression. We detected a strong reduction in CD63-phluorin signal in the CNS blood vessels on 3 dpf (Fig. 2E), as well as in the midbrain ventricle of 4 dpf *vps4a^T248I^* mutants (Figs. 2F,G). We quantified the CD63-pHluorin signal in the midbrain ventricle and normalized to background signal from the surrounding CNS tissue. The relationship between CD63-pHluorin signal in the midbrain ventricle is moderately correlated to signal in the muscle cells in wild-type siblings but not in mutants (Fig. 2H). We then calculated a ratio of the normalized pixel intensity of the midbrain ventricle CD63-pHluorin signal to the muscle cell CD63-pHlourin signal and found that the ventricle/muscle cell intensity ratio is significantly decreased in *vps4a^T248I^*mutants compared to wild-type siblings (*p* = 0.0382 (exact), nWT = 16, n*vps4a(T248I)* = 8, Mann-Whitney test) (Fig. 2I). This result indicates that there is a decrease in the number of exosomes present in the brain ventricle of mutants, supporting the hypothesis that biogenesis of these extracellular vesicles is reduced by the *vps4a^T248I^* mutation.

### Transcriptional changes in the *vps4a^T248I^* mutant

Considering the broad changes in endosomal morphology and the reduction in circulating exosomes in the CNS of *vps4a^T248I^* mutants, we next examined the gross morphology of the brain. Using whole mount immunohistochemistry, we labelled the axonal fibers of mutants and siblings with acetylated tubulin antibody to visualize axon morphology. No obvious changes in gross morphology between wild-type siblings and *vps4a^T248I^* mutants were apparent (Fig. 3A). The lack of obvious developmental defects is likely due to the presence of maternal mRNA. However, we noted the presence of pyknotic nuclei (Fig. 3B), which are associated with cell death (Hou et al., 2016). We quantified the number of these abnormal cells and found that *vps4a^T248I^* mutants have significantly more abnormal cells in the CNS compared to wild-type siblings and that cell death increases during development (5 dpf, *p* < 0.0001 (exact), nWT = 12, n*vps4a(T248I)* = 14, Mann-Whitney Test; 7 dpf, *p* < 0.0002 (exact), nWT = 8, n*vps4a(T248I)* = 7, Mann-Whitney Test).

**Figure 3.**
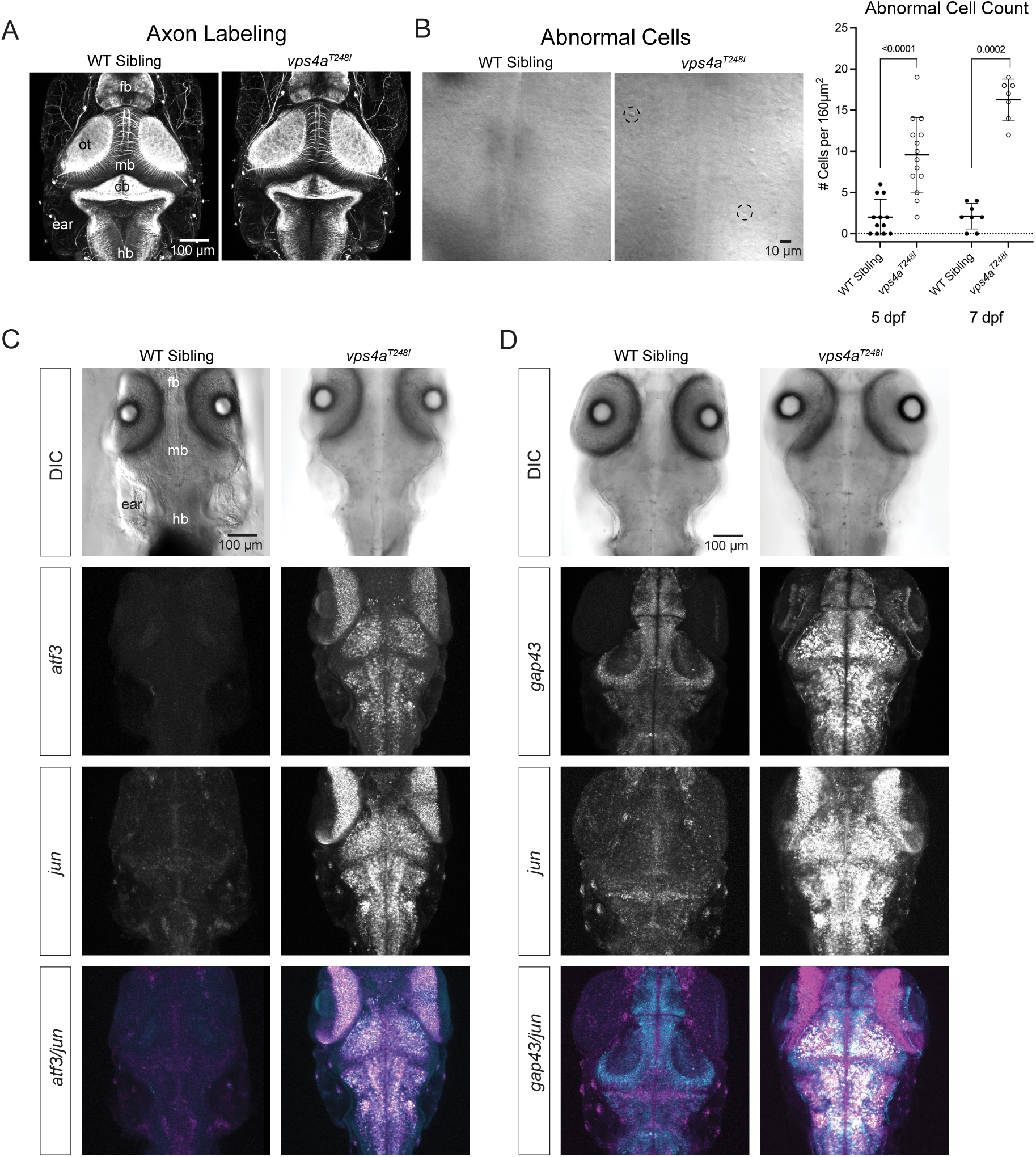
*vps4a^T248I^*mutants upregulate stress response and regeneration genes in the CNS. **A,** Axons labeled with acetylated tubulin antibody staining of *vps4a^T248I^* mutants show no gross morphological defects. Fb, forebrain; ot, optic tectum; mb, midbrain; cb, cerebellum; hb, hindbrain. **B,** *vps4a^T248I^* mutants have an increase of morphologically abnormal, pyknotic cells in the CNS. Two examples are highlighted with black dashed circles. The number of abnormal cells in the *vps4a^T248I^* mutants was quantified and found to be significantly higher than in wild-type siblings at both 5 dpf and 7 dpf; Mann-Whitney test, 5 dpf, *p* < 0.0001 (exact), nWT = 12, n*vps4a(T248I)* = 14; 7 dpf, *p* < 0.0002 (exact), nWT = 8, n*vps4a(T248I)* = 7. **C,** Maximum intensity projections of *atf3* and *jun* mRNA expression in the hindbrain of 5 dpf wild-type sibling and *vps4a^T248I^* mutants. **D,** Maximum intensity projections of *gap43* and *jun* mRNA expression in the hindbrain of 5 dpf wild-type sibling and *vps4a^T248I^* mutants.

As exosomes carry cargo containing microRNA and DNA and have been implicated in the regulation of DNA transcription (Kalluri and LeBleu, 2020), we assayed for transcriptional changes in the *vps4a^T248I^* mutant by performing a differential gene expression analysis on a bulk RNAseq data set that was initially used to find the mutation. We then used HCR RNA-FISH to validate the top log2 fold changes in gene expression. We found that the immediate early response genes *atf3* and *jun,* and the axonal regeneration gene *gap43* were dramatically upregulated in mid- and hindbrain regions in the *vps4a^T248I^*larvae (Fig. 3C, D). These striking changes in gene expression are consistent with a failure in intercellular communication by exosomes, and/or endosomal processing within neurons.

### Impaired vision in *vps4a^T248I^* mutants

The clinical profile of human patients with missense mutations in *VPS4A* includes visual deficits with a range of symptoms such as retinal dystrophy and cataract formation (Rodger et al., 2020; Seu et al., 2020). Although we did not observe expanded melanophores in the *vps4a^T248I^* mutant, which are indicative of blindness in larvae, we did find pronounced changes in visual function and gene expression in the retina. The optomotor response (OMR) in zebrafish is a reflex that can be evoked by a visual stimulus such as moving stripes, which causes the fish to swim in the direction of perceived motion, resulting in stabilization relative to the environment (Kist and Portugues, 2019). Compared to wild-type siblings, *vps4a^T248I^* mutant larvae have a greatly reduced OMR, suggesting they are blind (*p* < 0.0001 (exact), nWT = 12, n*vps4a(T248I)* = 10, Mann-Whitney test) (Fig. 4A). To investigate this defect further, we recorded electroretinograms (ERG) to assess the light induced changes in the electrical potential of the whole retina. The b-wave amplitude generated from an ERG can provide insight into the function of the outer retina (Niklaus et al., 2017). A range of light intensities (from the dimmest at log-4, to the brightest at log0) were tested and we found that the b-wave amplitude in *vps4a^T248I^* mutants is significantly decreased compared to wild-type siblings (log 0, *p* < 0.0001, nWT = 23, n*vps4a(T248I)* = 34; log -1, *p* < 0.0001, nWT = 23, n*vps4a(T248I)* = 33; log -2, *p* < 0.0001, nWT = 23, n*vps4a(T248I)* = 34; log -3, *p* = 0.0014, nWT = 23, n*vps4a(T248I)* = 34; log -4, *p* = 0.1989, nWT = 23, n*vps4a(T248I)* = 34, Kruskal-Wallis test) (Fig. 4B). In addition, we found that *jun* and *gap43* are upregulated in the retina of *vps4a^T248I^* mutants, specifically in the photoreceptor and retinal ganglion layers (Fig. 4C). We also detected reduced *opn1sw1* expression in the photoreceptors of mutants. Together, these analyses indicate that the *vps4a^T248I^* mutants have pronounced visual defects originating in the retina. Notably, the decrease in b-wave amplitude is variable among individual mutants and may not fully explain the absence of the OMR.

**Figure 4.**
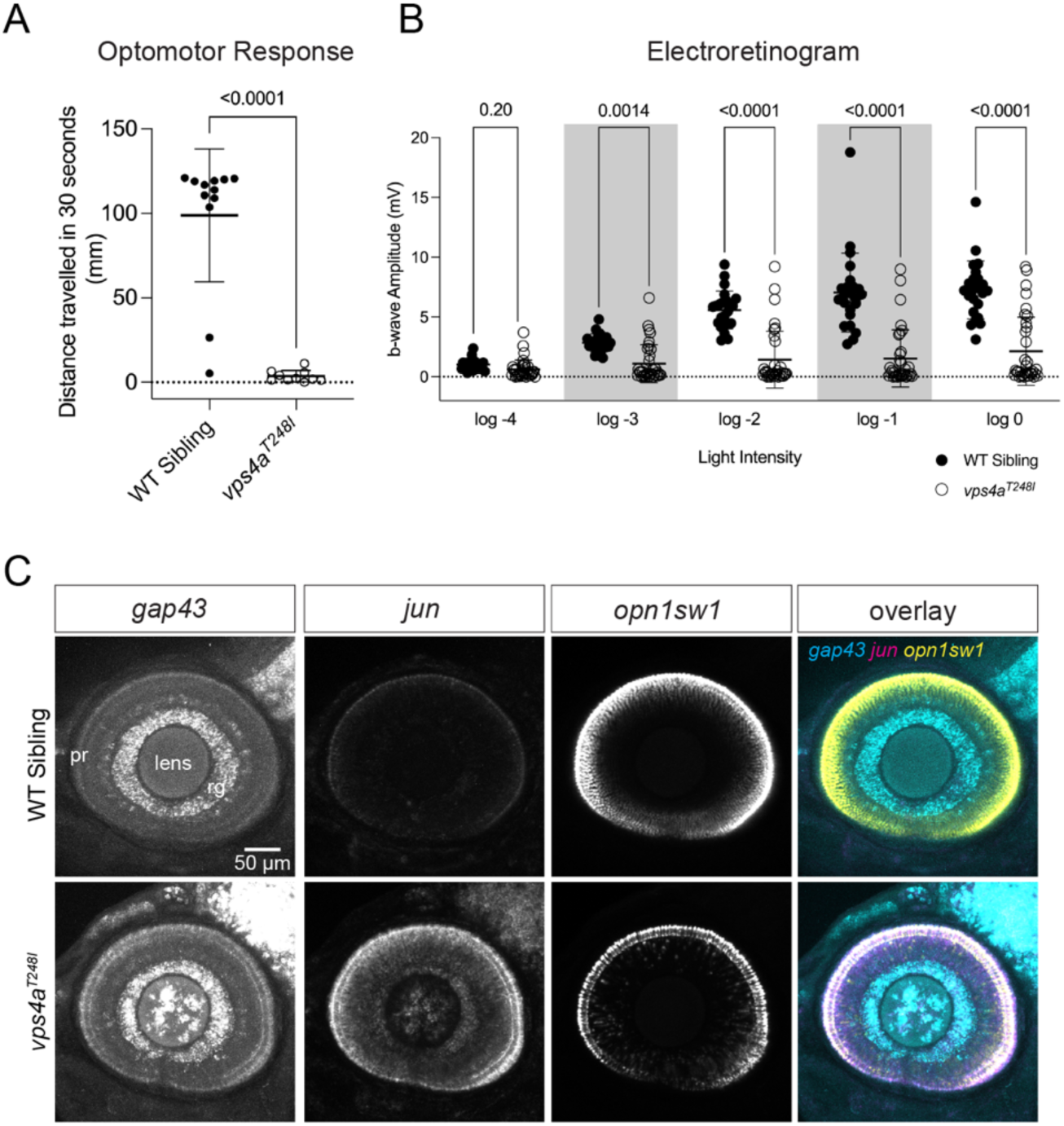
Impaired vision in *vps4a^T248I^* mutants. **A,** 5 dpf *vps4a^T248I^* mutants (N = 10) have a reduced optomotor response compared to wild-type siblings (N = 12); Mann-Whitney test, *p* < 0.0001 (exact). **B,** Decreased b wave amplitude in electroretinograms of 5-6 dpf *vps4a^T248I^* mutants (N = 34) in comparison to wild-type siblings (N = 23) at brighter light intensities (log -4 to log 0); Kruskal-Wallis test (log 0, *p* < 0.0001; log -1, *p* < 0.0001; log -2, *p* < 0.0001; log -3, *p* = 0.0014; log -4, *p* = 0.1989). **C,** HCR RNA-FISH of *gap43*, *jun* and *opn1sw1* in the eye of *vps4a^T248I^* mutants compared to wild-type siblings. Pr, photoreceptors; Rg, retinal ganglion cells, lens.

### Selective vestibular defects in *vps4a^T248I^* mutants

At the free-swimming stage of development (5 dpf), *vps4a^T248I^*larvae do not maintain an upright posture, swim sideways or upside down, and do not inflate their swim bladder, which is classically associated with functional deficits in vestibular hair-cells (Nicolson, 2017). To further investigate this phenotype in *vps4a^T248I^*mutant larvae, we tested vestibular function by assessing vestibular induced eye movements (VIEM) (Mo et al., 2010). Larvae with loss of hair-cell function mutations have reduced eye movements in response to head rotation (Nicolson, 2017), however, we did not observe a significant difference between the *vps4a^T248I^*mutants and wild-type siblings (*p* = 0.4632 (exact), nWT = 18, n*vps4a(T248I)* = 16, Mann-Whitney test) (Fig. 5A).

**Figure 5.**
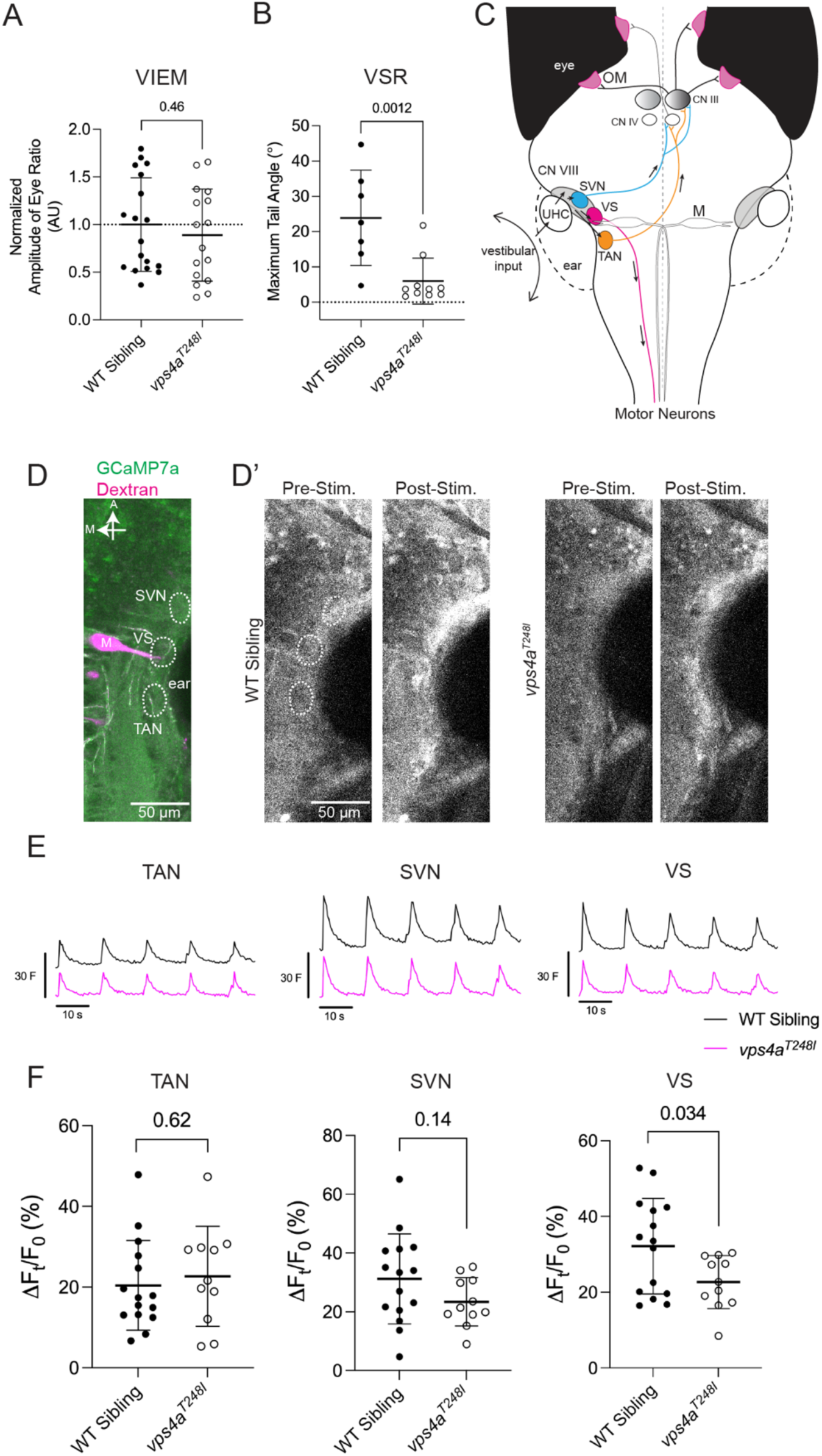
*vps4a^T248I^*mutants have selective defects in vestibular circuitry. **A,** 5 dpf *vps4a^T248I^* mutants (N = 16) do not have a significantly different VIEM compared to wild-type siblings (N = 18); Mann-Whitney text, *p* = 0.4632 (exact). **B,** 5 dpf *vps4a^T248I^* mutants (N = 10) have a reduced VSR compared to wild-type siblings (N = 7); Mann Whitney test, *p* = 0.0012 (exact). **C,** Simplified schematic showing the flow of information evoked by head rotation from the inner ear to hindbrain nuclei, vestibulo-ocular neurons, and motor neurons. UHC, utricular hair cells; CN VIII, VIIIth nerve; SVN, superior vestibular nucleus; VS, vestibulospinal cells; TAN, tangential neurons; M, Mauthner cell; CN IV, trochlear nucleus; CN III, oculomotor nucleus; OM, oculomotor muscles. **D,** 5 dpf larva showing basal GCamP7a expression and backfill injected Rhodamine-Dextran marking the Mauthner cell. White dotted lines mark the TAN, SVN and VS. **D’** Pre- and post-stimulus GCaMP7a levels in vestibular region in a representative wild-type sibling and *vps4a^T248I^* mutant. The TAN, SVN and VS ROIs are outlined by dotted lines. **E,** Average trace of normalized fluorescence (F) for the TAN, SVN and VS of wild-type siblings and *vps4a^T248I^* mutants during auditory stimulation at 600Hz. **F,** Percentage change in fluorescence of the TAN, SVN and VS of wild-type siblings and *vps4a^T248I^* mutants during auditory stimulation at 600Hz. The VS in *vps4a^T248I^* mutants (N = 11) have a reduced change in fluorescence compared to wild-type siblings (N = 15); unpaired t test, *p* = 0.0337 (exact), *t* = 2.252, df = 24. There is no significant difference between mutants and wild-type siblings in the TAN (unpaired t test, *p* = 0.1405 (exact), *t* = 1.524, df = 24) or SVN (unpaired t test, *p* = 0.6224 (exact), *t* = 0.4988, df = 24).

As mutant larvae have pronounced postural defects, we also tested the vestibulospinal reflex (VSR), which assesses vestibular induced motor function in the trunk (Gao and Nicolson, 2021). We compared the maximum tail angle between wild-type siblings and mutants and found the *vps4a^T248I^* mutants had a significantly reduced tail angle (*p* = 0.0012 (exact), nWT = 7, n*vps4a(T248I)* = 10, Mann-Whitney test) (Fig. 5B). The normal VIEM along with the reduced VSR result suggests that *vps4a^T248I^* mutants have selective defects in vestibular pathways.

In zebrafish, acceleration of the head is detected by utricular hair cells (Tanimoto et al., 2009; Mo et al., 2010; Sun et al., 2018), which then transmit signals to the utricular afferents of the VIIIth cranial nerve (Fig. 5C). The VIIIth nerve innervates the superior vestibular nucleus (SVN), vestibulospinal cell (VS) and tangential nucleus (TAN), as well as the Mauthner cell in the hindbrain (Bianco et al., 2012; Jia and Bagnall, 2022; Liu et al., 2022; Hamling et al., 2023b). Signals from the SVN and TAN are relayed by the oculomotor and trochlear nuclei to oculomotor muscles to produce eye movements associated with the VIEM (Fig. 5C;(Bianco et al., 2012; Liu et al., 2022; Sugioka et al., 2023)). VS neurons directly innervate motor neurons in the trunk to produce tail movements (Liu et al., 2022; Sugioka et al., 2023). To assess the function of the VSR circuit, we used a pan-neuronal GCaMP7a calcium indicator to visualize neuronal activity (Muto et al., 2013). Recently it was demonstrated that utricular hair cells in zebrafish larvae respond to high frequency tones, analogous to the responses of vestibular hair cells to bone-conducted vibration in mammals (Privat et al., 2019; Tanimoto et al., 2022). To stimulate the vestibular circuitry mediating the VIEM and VSR, we delivered pulses of 600 Hz at 110 dB using a mini-speaker attached to the glass slide on which larvae were mounted for calcium imaging. Electron microscopy data has shown that the lateral dendrite of the Mauthner cell is adjacent to the VS nucleus (Liu et al., 2022). To visualize this dendrite, we performed backfill injections of rhodamine dextran prior to calcium imaging, enabling us to use the Mauthner cell as a landmark for the approximate locations of the SVN, VS and TAN (Fig. 5D). Upon tone stimulation, neurons in all three nuclei responded with an increase in fluorescence (Fig. 5D’). Nevertheless, the average raw fluorescence trace for the VS region has a slightly lower peak amplitude in the *vps4a^T248I^*mutant compared to wild-type siblings (Fig. 5E, right).

Analysis of the group data from individual fish indicates that the percentage change in normalized fluorescence in the VS region is reduced in *vps4a^T248I^* mutants (*p* = 0.0337 (exact), nWT = 11, n*vps4a(T248I)* = 15, *t* = 2.252, df = 24)(Fig. 5F, right). As predicted by our behavioral analysis, we did not observe a significant difference in the raw fluorescence trace or percentage change in normalized fluorescence in the TAN or SVN between *vps4a^T248I^* mutants and wild-type siblings (TAN *p* = 0.6224 (exact), SVN *p* = 0.1405 (exact), nWT = 11, n*vps4a(T248I)* = 15, *t =* 0.4988 (SVN), df = 24 (SVN), *t* = 1.524 (TAN), df = 24 (TAN)) (Figs. 5E,F, left and center).

Overall, our calcium imaging data confirm that the VIEM circuit is intact and suggest that reduced activity in the VS neurons, which directly innervate motor neurons, contributes in part to the reduction in tail movements in response to vestibular cues in the *vps4a^T248I^* mutants.

### Intact ascending auditory pathway in *vps4a^T248I^* mutants

In zebrafish, auditory signals excite saccular hair cells, which are innervated by the afferent neurons of the statoacoustic ganglion (or VIIIth nerve). If the auditory cues are sufficiently loud enough to startle, the signal is transmitted to the giant escape fiber neuron known as the Mauthner cell, which directly activates motor neurons (Kohashi and Oda, 2008; Sillar, 2009; López-Schier, 2019). Less intense sounds activate hindbrain neurons within the medial octavolateralis nuclei region, which then communicate with the torus semicircularis for further auditory processing (Echteler, 1984; Tanimoto et al., 2009; Inoue et al., 2013; Vanwalleghem et al., 2017) (Fig. 6A). *vps4a^T248I^*mutant larvae are largely unresponsive to tapping on the petri dish, suggesting that hearing is profoundly impaired. We assessed the severity of this defect by quantifying the auditory evoked behavioral response (AEBR), which is a startle response to loud, pure tone stimuli. *vps4a^T248I^* mutant larvae responded to pure tones significantly less compared to wild-type siblings, indicating that mutants have pronounced defects (*p* < 0.0001 (exact), nWT = 15, n*vps4a(T248I)* = 20, Mann-Whitney test) (Fig. 6B). The pan-neuronal GCaMP7a calcium indicator was used to examine neuronal activity in the saccular hair cells and statoacoustic ganglion in response to the same 110 dB 600 Hz stimulus (Figs. 6C,D). In saccular hair cells, we observed robust responses in average raw fluorescence traces (Fig. 6E) and comparable calcium transients in wild-type or mutant individual fish (*p* = 0.4162 (exact), nWT = 20, n*vps4a(T248I)* = 13, *t* = 0.8228, df = 35) (Fig. 6F). Comparable responses were also seen in the statoacoustic ganglion (*p* = 0.8748 (exact), nWT = 20, n*vps4a(T248I)* = 13, *t* = 0.1588, df = 31) (Figs. 6E,F). These results are consistent with LysoTracker labeling, which shows no significant difference in dye uptake between wild-type siblings and *vps4a^T248I^*mutant in hair cells in the saccule (*p* = 0.0934 (exact), nWT = 6, nvps4a(T248I) = 9, *t* = 1.810, df = 13), utricle (*p* = 0.6314 (exact), nWT = 6, nvps4a(T248I) = 9, *t* = 0.4913, df = 13) or anterior cristae (*p* = 0.3233 (exact), nWT = 10, nvps4a(T248I) = 10, *t* = 1.016, df = 18).

**Figure 6.**
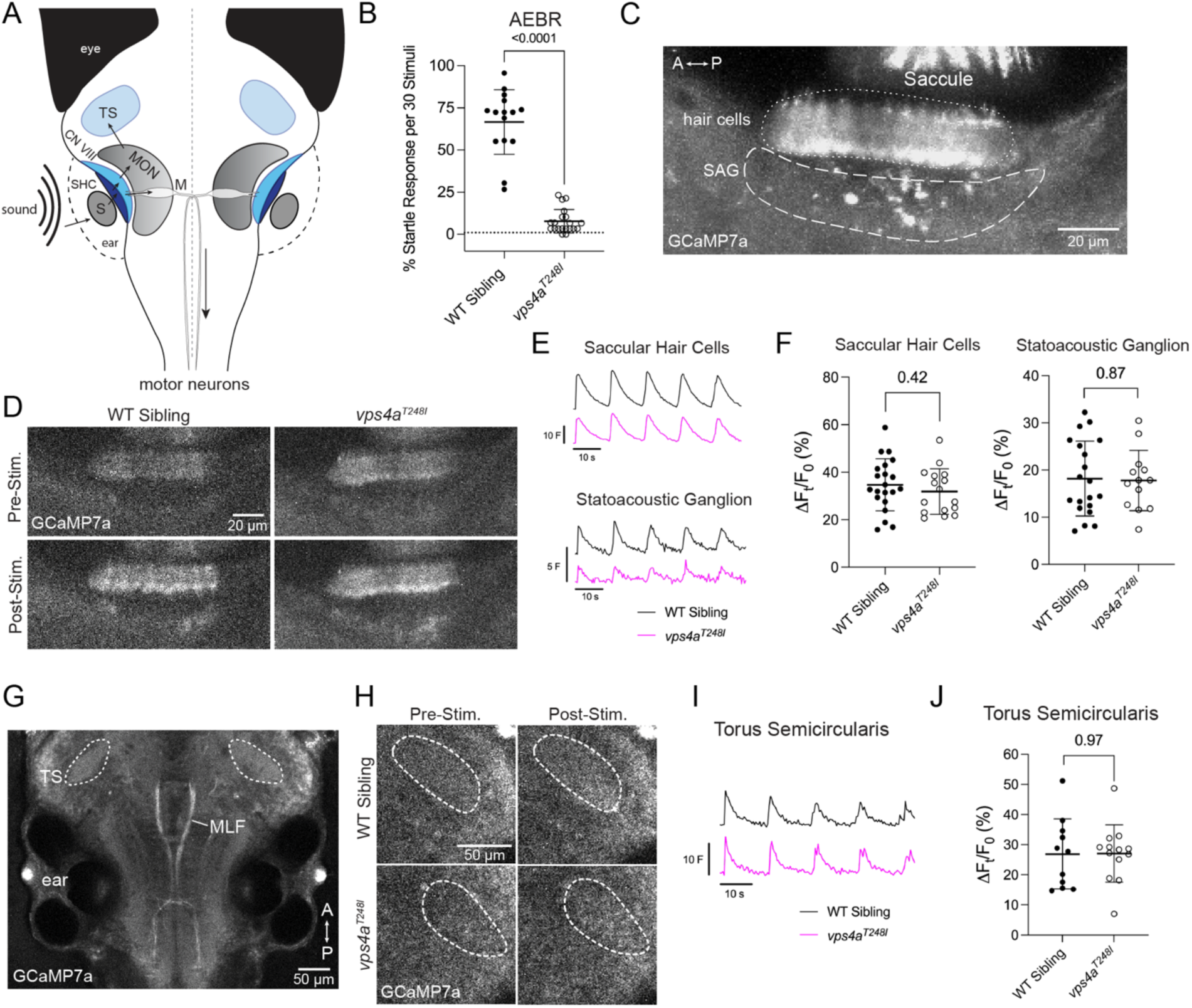
Activation of the ascending auditory pathway in *vps4a^T248I^* mutants. **A,** Simplified schematic showing the flow of information from an auditory stimulus to motor neurons or to the torus semicircularis. S, saccular otolith; SHC, saccular hair cells; CN VIII, VIIIth nerve; MON, medial octavolateralis nucleus; M, Mauthner cell; TS, torus semicircularis. **B,** 5 dpf *vps4a^T248I^* mutants (N = 20) have a greatly reduced AEBR compared to wild-type siblings (N = 15); Mann-Whitney test, *p* < 0.0001 (exact). **C,** Basal levels of GCaMP7a expression in the inner ear of 5 dpf larvae, focusing on the saccular hair cells (dotted line) and statoacoustic ganglion (SAG) (dashed line). **D,** Pre- and post-stimulus GCaMP7a levels in saccular hair cells and SAG in a representative wild-type sibling and *vps4a^T248I^* mutant larvae. **E,** Average traces of normalized fluorescence (F) for the saccular hair cells and SAG of wild-type siblings and *vps4a^T248I^* mutants during auditory stimulation at 600Hz. **F,** Percentage change in fluorescence of the saccular hair cells and SAG of wild-type siblings and *vps4a^T248I^* mutants during auditory stimulation at 600Hz. There is no significant difference between mutants (N = 13) and wild-type siblings (N = 20) in the saccular hair cells (unpaired t test, *p* = 0.4162 (exact), *t* = 0.8228, df = 35) or the SAG (unpaired t test, *p* = 0.8748 (exact), *t* = 0.1588, df = 31). **G,** Representative image of 5 dpf zebrafish basal levels of GCaMP7a expression in the torus semicircularis (TS). **H,** Pre-and post-stimulus GCaMP7a levels in the TS in representative wild-type sibling and *vps4a^T248I^*mutant larvae. **I,** Average trace of normalized fluorescence (F) for the TS of wild-type siblings and *vps4a^T248I^* mutants during auditory stimulation at 600Hz. **J,** There is no significant difference in percentage change fluorescence between *vps4a^T248I^* mutants (N = 13) and wild-type siblings (N = 11) in the TS; unpaired t test, *p* = 0.9672 (exact), *t* = 0.04157, df = 22.

The auditory circuitry within the hindbrain for non-startle responses is less well defined in larvae, therefore we focused on a midbrain region of the ascending auditory pathway and examined neuronal activity in the torus semicircularis, which is akin to the inferior colliculus in mammals and key for auditory processing (McCormick, 1999; Edds-Walton and Fay, 2008; Vanwalleghem et al., 2017). This midbrain structure is located beneath the optic tectum at the level of the semicircular canals and saccule of the inner ear and the medial longitudinal fasciculus (MLF) (Fig. 6G). We observed a visible increase in GCaMP7a reporter activity in the torus semicircularis of both wild-type siblings and *vps4a^T248I^*mutants after pure tone stimulation (Fig. 6H). No significant differences were detected in the average raw fluorescence traces or percentage change in normalized fluorescence between wild-type siblings and *vps4a^T248I^* mutants for the torus semicircularis (*p* = 0.9672 (exact), nWT = 11, n*vps4a(T248I)* = 13, *t* = 0.04157, df = 22) (Figs. 6I,J). Together, these data indicate that the ascending auditory pathway of *vps4a^T248I^* mutants including the inner ear and central circuitry is functional.

### Defective sensorimotor transformation in *vps4a^T248I^* mutants

Although *vps4a^T248I^* mutants have a greatly reduced acoustic startle response, our data demonstrate that auditory cues activate the ascending auditory pathway. Notably, mutant larvae are not paralyzed, however, they are largely inactive at the free-swimming stage. Therefore, we assessed whether *vps4a^T248I^*mutants have a general motor system defect that prevents motor responses to sensory cues.

To test whether startle reflexes are globally affected, we measured the ability of *vps4a^T248I^* mutants to produce a deep C-bend of the trunk in response to contact of a stiff probe with the dorsal region of the head. We observed that both wild-type siblings and *vps4a^T248I^* mutants displayed robust C-bend motions of their trunks in response to touch (Figs. 7A,B; *p* = 0.1079 (exact), nWT = 10, n*vps4a(T248I)* = 11, *t* = 1.687, df = 19). This result suggests that touch receptors, along with Mauthner cells and other vital components of motor output such as spinal cord neurons, motor neurons, and muscle cells are functional. In addition, axonal fibers in superficial layers of the skin, the hindbrain, and anterior trunk regions did not show any changes in gross morphology in *vps4a^T248I^* mutants (Fig. 7C). These data point to a deficit within central circuitry in *vps4a^T248I^* mutants as downstream motor components of the startle reflex are unaffected.

**Figure 7.**
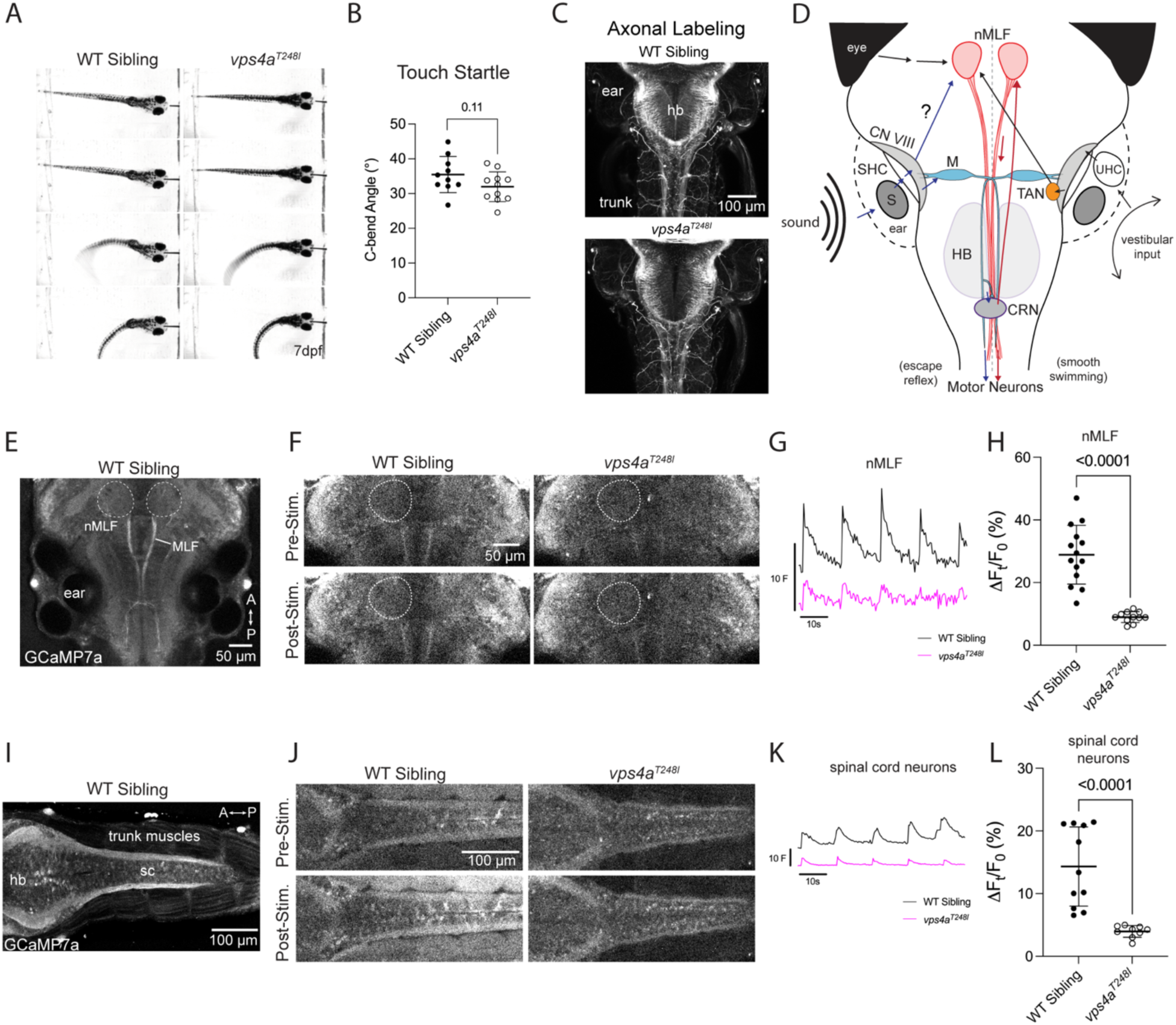
*vps4a^T248I^*mutants display defects in sensorimotor transformation. **A,** 7 dpf *vps4a^T248I^* mutant larvae have an intact startle reflex when touched. **B,** 7 dpf *vps4a^T248I^* mutants (N = 11) do not have a significantly different C-bend angle compared to wild-type siblings (N = 10); unpaired t test, *p* = 0.1079 (exact), *t* = 1.687, df = 19. **C,** Acetylated tubulin antibody staining of *vps4a^T248I^* mutants shows no gross morphological defects in hindbrain/spinal cord or somatosensory axons. **D,** Simplified schematic showing the flow of information from a tone stimulus from the inner ear to motor neurons via Mauthner cell and nMLF pathways. S, saccular otolith; SHC, saccular hair cells; CN VIII, VIIIth nerve; M, Mauthner cell; HB, hindbrain; CRN, cranial relay neurons; nMLF, nucleus of the medial longitudinal fasciculus. **E,** Representative image of 5 dpf zebrafish, showing location of nMLF expressing basal levels of GCaMP7a. **F,** Pre- and post-stimulus GCaMP7a levels in the nMLF in a representative wild-type sibling and *vps4a^T248I^* mutant. **G,** Average trace of normalized fluorescence (F) for the nMLF of wild-type siblings and *vps4a^T248I^*mutants during tone stimulation at 600Hz. **H,** *vps4a^T248I^* mutants (N = 11) have a significantly reduced percentage change fluorescence in the nMLF compared to wild-type siblings (N = 14); unpaired t test, *p* < 0.0001 (exact), *t* = 6.937, df = 23. **I,** Representative image of 5 dpf zebrafish trunk showing spinal cord neurons expressing basal levels of GCaMP7a. **J,** Pre-and post-stimulus GCaMP7a levels in the spinal cord motor neurons of representative wild-type sibling and *vps4a^T248I^*mutant. **K,** Average trace of normalized fluorescence (F) for the spinal cord motor neurons of wild-type siblings and *vps4a^T248I^* mutants during tone stimulation at 450Hz. **L,** *vps4a^T248I^*mutants (N = 9) have a significantly reduced percentage change fluorescence in spinal cord neurons compared to wild-type siblings (N = 11), Mann-Whitney test, *p* < 0.0001 (exact).

As described above, auditory signals from the statoacoustic ganglion are directly transmitted to the Mauthner cell, which innervates trunk motor neurons to generate an escape reflex. However, the 110 dB intensity of our tone stimulus did not result in detectable activation of the Mauthner cell soma, which is consistent with a previous imaging study (Marsden and Granato, 2015). Instead, we observed a reliable response in the midbrain nucleus of the medial longitudinal fasciculus (nMLF) (Figs. 7D, E-H). The nMLF is a vital component of the descending motor system, serving as a command center for forward locomotion (Sankrithi and O’Malley, 2010; Koyama et al., 2011; Severi et al., 2014; Thiele et al., 2014; Wang and McLean, 2014; Migault et al., 2018; Shimazaki et al., 2019; Xu et al., 2021). The nMLF contains subpopulations of vGlut1 and vGlut2 positive neurons, which control both fast escape-swimming behaviors as well as slow forward locomotion (Severi et al., 2014; Berg et al., 2023). Neurons in the nMLF were recently shown to respond to hindbrain cranial relay neurons (CRN; Fig. 7D), which can be activated by aversive auditory stimulation of Mauthner cells, leading to smooth swimming after an initial C-bend response (Xu et al., 2021). Although the auditory circuitry to the nMLF activated by non-startle stimuli has not been characterized, the nMLF has been implicated in fine postural control via inputs from the TAN of the ascending vestibuloocular pathway (Sugioka et al., 2023). Notably, activation of neurons in the TAN in *vps4a^T248I^* mutants is comparable to wild-type responses.

Considering that the nMLF is central to motor function in response to sensory cues, we compared GCaMP7a signals in this cluster of neurons between wild-type siblings and *vps4a^T248I^* mutants in response to the 600 Hz stimulus. These midbrain clusters can be identified based on their prominent axonal tracts that descend near the midline (Fig. 7E). We found that the peak amplitudes of calcium transients in the average raw fluorescence trace in the mutant were markedly decreased (Figs. 7F-G). In addition, the percentage change in normalized fluorescence in the nMLF was significantly reduced in all individual *vps4a^T248I^* mutants tested (*p* < 0.0001 (exact), nWT = 14, n*vps4a(T248I)* = 11, *t* = 6.937, df = 23) (Fig. 7H).

We next investigated the neuronal activity of the spinal cord in response to tone stimulation using the pan neuronal GCaMP7a reporter (Fig. 7I). The average raw fluorescence trace of the *vps4a^T248I^* mutants show lower peaks in response to tone stimulation (Fig. 7K). In the examples shown in Figure 8J, a second reporter (GCaMP7a expressed under a muscle cell promoter, (Leung et al., 2019)) in the trunk muscles shows activation in the wild-type larva but none in the mutant. Overall, the percentage change in normalized fluorescence is strongly reduced in *vps4a^T248I^* mutants (*p* < 0.0001 (exact), nWT = 11, n*vps4a(T248I)* = 9, Mann-Whitney test) (Fig. 7L). Together, our results indicate that tone stimulation fails to trigger responses in the nMLF descending motor pathway and downstream trunk motor neurons in *vps4a^T248I^*mutants.

## Discussion

Our study identifies a role for *vps4a* in sensorimotor transformation during larval development in zebrafish. The T248I mutation in *vps4a,* which was isolated in a forward genetic screen for balance and hearing deficits, profoundly affects enzymatic function, leading to defects in ESCRT filament disassembly and membrane scission-dependent processes such as exosome biogenesis. In *vps4a^T248I^*larvae, we observe an increase in the expression of *jun* and *atf3* stress response genes and a progressive loss in cellular homeostasis in the developing brain and retina. Although identified as a deafness mutant, the function of the inner ear in *vps4a^T248I^* larvae is unaffected. Combined behavioral analyses and *in vivo* imaging of circuit activity demonstrate that the ascending auditory pathway and descending motor pathways are intact in *vps4a^T248I^* mutants, however, mutants do not respond to aversive acoustic stimuli. These data suggest that loss of *vps4a* function induces early onset of deficits in the transformation of sensory signals, particularly auditory, vestibular, and visual input into motor responses.

### Defective disassembly of ESCRT filaments and decreased exosome biogenesis in zebrafish *vps4a* mutants

Our biochemical analyses indicate that the substitution of the corresponding threonine residue with isoleucine in the ATPase domain of yeast *Vps4* strongly reduces ATPase activity and ESCRT-III filament disassembly. *In vivo* imaging of the CNS revealed enlargement of acidic compartments in larval mutants, which is consistent with previous studies in yeast *vps4* mutants and human *VPS4A* patient fibroblasts (Bishop and Woodman, 2000; Frankel et al., 2017; González et al., 2017; Willén et al., 2017). Enlarged endosomal compartments are likely due to the disruption of endosomal processing and when membrane fails to invaginate and form intraluminal vesicles via the ESCRT-mediated scission process. These vesicles can be released from nearly all cell types as exosomes and contain a variety of signaling molecules, such as proteins, lipids, and nucleic acids (Colombo et al., 2014; Zhang et al., 2019; Gurunathan et al., 2021; Waqas et al., 2022). Exosomes have been implicated in a several diseases such as cancers and neurodegenerative diseases due to the vital role they play in intercellular communication (Lim and Lee, 2017; Bebelman et al., 2020; Dilsiz, 2020; Gomes et al., 2020; Duarte-Silva et al., 2022; Han et al., 2023). A previous study on erythroid cells, which found that transferrin receptor trafficking, a process dependent on proper endosomal sorting and exosome secretion, is disrupted in human patients with *de novo VPS4A* mutations (Seu et al., 2020). Here we show direct evidence for reduced exosome biogenesis using a CD63-pHluorin marker of circulating exosomes. Both in blood vessels and brain ventricles, *vps4a^T248I^* mutants show a marked decrease in CD63-pHlourin signal. Disruptions to intercellular signaling through exosomes can manifest as transcriptional changes (Dobrowolski and De Robertis, 2012; Budnik et al., 2016; Gurung et al., 2021). We found that several genes associated with transcription (*atf3* and *jun*), or regeneration (*gap43*) were highly upregulated in the CNS of *vps4a^T248I^*mutants. *atf3* and *jun* are known stress response genes, whereas upregulation of *gap43* may be a compensatory response to disruption of axonal processes. Although exosomes mediate cell-to-cell signaling and are important for cellular homeostasis, it is unclear whether the transcriptional changes in *vps4a^T248I^*mutants are due to (*i*) reduced exosome signaling, (*ii*) disruptions in endosomal trafficking, or (*iii*) a combination of both defects. Alternatively, other disruptions to membrane scission in the CNS may be involved as well. The expression of stress response genes in the CNS in zebrafish mutants during larval development is consistent with the microcephaly seen in human patients. While we did not observe changes in gross morphology of the CNS, cell death in the brain was evident and increased during development. We speculate that more pronounced defects would be evident in older animals, however, we limited our analyses to early stages of larval development during which we could identify early onset defects.

### Sensory and motor deficits

Patients with *VPS4A* mutations are reported to have sensory deficits relating to vision including retinal dystrophy and cataracts (Rodger et al., 2020). With respect to visual function in *vps4a^T248I^* mutants, larvae fail to swim in response to moving visual cues and outer retinal responses to light are significantly reduced, signifying defects in vision that align with transcriptional changes in the retina. The reduction in outer retinal function was variable among mutants, whereas the optomotor response was severely reduced in all *vps4a^T248I^* mutants, suggesting that loss of photoreceptor function provides only a partial explanation of the absence of vision-associated motor reflexes. Notably, human patients are reported to have difficulties in fixing and following a moving object, which may be due to poor visual acuity or a combination of poor vision and oculomotor pathway dysfunction.

Loss of central control of muscle tone, ataxia and motor delays are common traits found in patients with neurodevelopmental disorders, including *VPS4A* patients (Rodger et al., 2020; Seu et al., 2020). *vps4a^T248I^*larvae share some of these features, such as lack of control over posture and sensorimotor deficits. Vestibulospinal reflexes, which result in corrective movements of the trunk, are greatly diminished in mutants. However, not all vestibular function in lost; *vps4a^T248I^*mutants still produce eye movements in response to rotation of the head.

Consistent with the behavioral data, neurons of the TAN and SVN, which are part of vestibuloocular pathway (Bianco et al., 2012; Liu et al., 2022), show comparable activity in the *vps4a^T248I^* mutant to wild-type neurons. This data indicates that perturbation of Vps4a activity does not result in global defects in vestibular function, or in brain function in general, but rather that specific circuits are more sensitive to mutations in *vps4a*. Normal vestibular induced eye movements and activation of TAN and SVN neurons also indicate that excitation of utricular hair cells and vestibular afferent neurons is unaffected in *vps4a^T248I^* mutants. In contrast to the vestibuloocular pathway, the vestibulospinal circuit is impaired when *vps4a* is mutated.

Hindbrain VS neurons, which directly innervate trunk motor neurons, have decreased neuronal activity in mutants. Recently it was shown that ablation of VS neurons leads to modest defects in posture in zebrafish larvae (Hamling et al., 2023a), suggesting that other vestibular circuits contribute to postural control. As such, the reduced activation of the vestibulospinal pathway in *vps4a^T248I^* mutants may only partially account for the inability to maintain an upright posture.

Further evidence for a loss in central control of motor responses emerged from our analysis of the absence of an acoustic startle reflex in *vps4a^T248I^*mutants. Despite the lack of response to aversive acoustic stimuli, *vps4a^T248I^*mutants have comparable activation of the ascending auditory pathway as seen in wild-type siblings. Deficits in the descending motor pathways or the spinal cord are also not the cause for the absence of the motor output in response to visual or tone stimulation. *vps4a^T248I^* mutants are not paralyzed and aversive touch stimuli activate escape responses, demonstrating that mutant larvae can generate robust tail movements. The above experiments indicate that both saccular afferent neurons and Mauthner cells are activated by sound and touch in *vps4a^T248I^* mutants, respectively, implying that a second pathway may be defective in generating acoustic startle reflexes. One such pathway may involve hindbrain spiral fiber neurons, which have been shown to be essential for Mauthner cell-dependent escape responses (Lacoste et al., 2015). Nevertheless, our *in vivo* imaging experiments using a non-startle tone stimulus revealed a pronounced reduction in the activity of midbrain nMLF neurons and downstream spinal cord neurons of *vps4a^T248I^* mutants. Neurons of the nMLF play a key role in receiving sensory input and sending motor commands to the spinal cord and are thought to play an analogous role to the interstitial nucleus of Cajal in the mammalian midbrain, which controls eye and head movements (Fukushima, 1987; Severi et al., 2014; Thiele et al., 2014). In zebrafish, both visual and vestibular systems provide sensory input to the nMLF (Bianco et al., 2012; Matsuda and Kubo, 2021; Liu and Bagnall, 2023).

Interestingly, TAN neurons of the vestibuloocular pathway, also play a dual role of controlling eye and body movements in zebrafish, and were recently shown to transmit signals about the roll axis of the body to the nMLF, implicating this circuit in fine control of posture (Sugioka et al., 2023). In *vps4a^T248I^* mutants, TAN neurons are also activated by the tone stimulus, yet postural defects are pronounced, further bolstering the notion that sensory input such as vestibular cues fail to trigger sensorimotor transformation within the nMLF.

In summary, our combined biochemical and *in vivo* analyses have revealed that loss of Vps4a function generates sensorimotor processing defects in the CNS of zebrafish larvae. Our study provides a better understanding of the cause of motor dysfunction when *vps4a* is mutated. Recent studies of exosome therapy for sensorimotor recovery after injury or resulting from acute or chronic neurodegenerative disease (Chen et al., 2020; Yari et al., 2022; Zhang et al., 2023) suggest that exosomes therapy may ameliorate the symptoms of neurodevelopmental disorders such as those associated with *VPS4A*.

## Acknowledgements

The authors would like to thank the past members of the Genetics Department of the Max Planck Institute in Tuebingen, Germany, particularly Tatjana Piotrowski and Stefan Schulte-Merker for their assistance in a small-scale allele screen. We would also like to thank Matthew Esqueda and Sivan Brodo-Abo for their help with animal care. This work was supported in part by a National Institute on Deafness and Other Communication Disorders award (DC0170046) and SICHL funding from the OHNS department of Stanford University to T.N.; a grant from the National Natural Science Foundation of China to S.S. (92254306); Swiss National Science Foundation (310030_204648) to S.N.

